# Differences in the Pupillary Responses to Evening Light between Children and Adolescents

**DOI:** 10.1101/2023.08.09.552691

**Authors:** Lauren E. Hartstein, Monique K. LeBourgeois, Mark T. Durniak, Raymond P. Najjar

## Abstract

**Purpose:** To assess differences in the pupillary light responses (PLRs) to blue and red evening lights between children and adolescents.

**Methods:** Forty healthy participants (8-9 years, n=21; 15-16 years, n=19) completed a PLR assessment 1 h before their habitual bedtime. After a 1 h dim-light adaptation period (<1 lux), baseline pupil diameter was measured in darkness for 30 s, followed by a 10 s exposure to 3.0×10^13^ photons/cm^2^/s of either red (627 nm) or blue (459 nm) light, and a 40 s recovery in darkness to assess pupillary re-dilation. Subsequently, participants underwent 7 min of dim-light re-adaptation followed by an exposure to the other light condition. Lights were counterbalanced across participants.

**Results:** Across both age groups, maximum pupil constriction was significantly greater (p< 0.001, η_p_^2^=0.48) and more sustained (p< 0.001, η_p_^2^=0.41) during exposure to blue compared to red light. For adolescents, the post-illumination pupillary response (PIPR), a hallmark of melanopsin function, was larger after blue compared with red light (p= 0.02, d=0.60). This difference was not observed in children. Across light exposures, children had larger phasic (p< 0.01, η_p_^2^=0.20) and maximal (p< 0.01, η_p_^2^=0.22) pupil constrictions compared to adolescents.

**Conclusions:** Blue light elicited a greater and more sustained pupillary response than red light across participants. However, the overall amplitude of the rod/cone-driven phasic response was greater in children than in adolescents. Our findings using the PLR highlight a higher sensitivity to evening light in children compared to adolescents, and continued maturation of the human non-visual photoreception/system throughout development.

## INTRODUCTION

Intrinsically photosensitive retinal ganglion cells (ipRGCs) are ocular photoreceptors that express the photopigment melanopsin and are maximally sensitive to short-wavelength blue light, with a peak sensitivity of ∼480nm ^1, 2^. ipRGCs integrate their intrinsic response to inputs from rods and cones, before transmitting the signal to non-visual centers in the brain such as the suprachiasmatic nucleus (SCN), the central timekeeper of bodily circadian rhythms^3, 4^, the olivary pretectal nucleus (OPN) which controls the pupillary light response (PLR)^5, 6^, and centers involved in sleep and alertness^7^.

The PLR offers a fast and reliable biomarker for testing the physiology and integrity of visual and non-visual photoreception. It generally consists of a rapid constriction of the pupil to an abrupt light stimulus. This transient constriction response is largely attributed to outer-retinal (rod/cone) activity^6^. If the light stimulation is maintained, the transient constriction is followed by a sustained constriction of the pupil (steady-state). The steady-state response consists of input from all photoreceptors; however, the ipRGC’s intrinsic contribution to this state is three-folds higher than long-wavelength (L-cones) and medium-wavelength (M-cones) cones^5^. After cessation of the light stimulus, signals from both the rods and ipRGCs contribute to the post-illumination pupil response (PIPR) in the first 1.7s, after which the PIPR is entirely driven by ipRGCs^8^. In adults, exposure to short-wavelength blue light results in faster, greater, and more sustained pupillary constriction than long-wavelength red light^9–11^. This more sustained response to blue light is consistent with greater stimulation of the ipRGCs. Therefore, components of the PLR can be used as an indicator of the ipRGC response and can be utilized as a marker in health and disease^12–14^.

Baseline pupil diameter increases throughout childhood, peaking in adolescence ^15–17^, then slowly decreases throughout adulthood ^18–20^. In adults age 21-70 years, a larger baseline pupil diameter was associated with a larger PIPR 6 s after exposure to blue light^18^. However, when controlling for baseline pupil diameter, age was not associated with PIPR, indicating that the inputs of the ipRGC to the pupillary pathway do not diminish with advancing age. Data on the maturation of the PLR throughout child development, however, remain limited. In premature infants (gestational age 38 weeks), the PLR is observed in response to blue but not red light^17^. However, the magnitude of the pupillary constriction, as well as baseline pupil size, is significantly smaller in premature infants than at 2 years of age^17^. In a longitudinal study of the PLR elicited by green light (530 nm) across children aged 6-24 months, baseline size, minimal pupil size, and maximum pupil constriction (% constriction with respect to baseline) increased with age, whereas latency (the time from light onset to the start of pupil constriction) decreased with age^21^. In a sample of children aged 6-17 years, the amplitude of pupillary constriction in response to green light increased until age 8, then plateaued^16^. PLR latency decreased across ages 6-9, then stabilized. Constriction and re-dilation time were not associated with age. Across children ages 5-15 years, the ipRGC-driven pupil response to light was observed to be robust and similar to those measured in adults^22^. Lastly, in another study examining the PLR between children aged 3-10 and 10-18 years, the younger group required a brighter red light stimulus to evoke a 5% pupil constriction than the older group^23^.

The circadian timing of a light exposure can influence the PLR^24^. In adults, circadian fluctuations in post-stimulus pupil size were observed in response to blue (463 nm) but not red (635 nm) light ^25^. Additionally, the ipRGC-driven pupil response in children is influenced by individual differences in 24-h light history^22^. Furthermore, across the lifespan, humans spend the vast majority of their time in indoor environments ^26, 27^, resulting in less exposure to bright light during the day and greater exposure to artificial light at night ^28^. Both the circadian timing of assessments and prior light history are largely unreported and uncontrolled for in previous studies of the PLR in children. The purpose of the present research was to (1) study the chromatic pupillary light reflex in response to red and blue evening light in children and adolescents while controlling for the confounding factors mentioned above; and (2) examine age-related differences across 2 distinct stages of child development.

## METHODS

### Participants

Children aged 8.0-9.9 years (n = 21, 12 female, M = 8.99 years, SD = 0.51 years) and adolescents aged 15.0-16.9 years (n = 19, 5 female, M = 15.78 years, SD = 0.55 years) completed a 6-day study protocol. Participants were recruited from the greater Boulder, CO, USA area. Prospective interested parents were screened through an online questionnaire and subsequent phone interview. Participants were excluded from the study for parent report of any of the following: clinical sleep disorders; behavioral problems; personal or family history of diagnosed narcolepsy, depression, psychosis, or bipolar disorder; pre-term delivery (< 35 weeks); low birth weight (< 5.5 lbs.); current use of caffeine, nicotine, alcohol, or medications affecting sleep, the circadian system, or light sensitivity; active allergies or asthma; developmental disabilities; neurological disorders; metabolic disorders; chronic medical conditions; head injury involving loss of consciousness; travel > 2 time zones in the 2 months before participation; sleep schedule varying > 2 h between weekdays and weekends; migraine or frequent headaches; eye disorders (excluding corrected refractive error) or color blindness (confirmed using the Ishihara Color Vision Test). All study procedures were approved by the University of Colorado Boulder Institutional Review Board. Parents provided written, informed consent and children/adolescents provided written assent to participate. Families were compensated for their participation.

### Protocol

Data collection occurred between June and October in 2021 and September through October in 2022. For 5 days, children followed a strict parent-selected sleep schedule (bedtime and wake time) with ≥ 9 h time in bed, whereas adolescents followed a self-selected sleep schedule of ≥ 8 h time in bed^29^. Adherence to the sleep schedule was confirmed through a wrist-worn actigraph (Spectrum Plus, Philips Respironics, Pittsburgh, PA, USA) and a daily sleep diary (parent-reported for the school-aged children and self-reported for the adolescents). Participants were instructed to avoid light exposure between bedtime and wake. Starting at wake time on Day 6, participants wore a pair of dark glasses (UVEX by Honeywell, Espresso lens) with an advertised visible light transmission of 15%^30^ in order to limit exposure to bright light and reduce individual variability in light history (**Fig. S1**). The transmittance of the glasses at 480nm was measured as 8.9%. Participants were instructed to remain indoors wearing the glasses throughout the day until beginning the dim-light adaptation at the laboratory that evening (**Fig. 1)**. Participants arrived at the lab 2.5 h before their scheduled bedtime for the pupillary assessment. They then entered a dimly-lit room (< 1 lux at angle of gaze) beginning 2 h before their scheduled bedtime and adapted to the dim-light for 1 h.

**Fig. 1:**
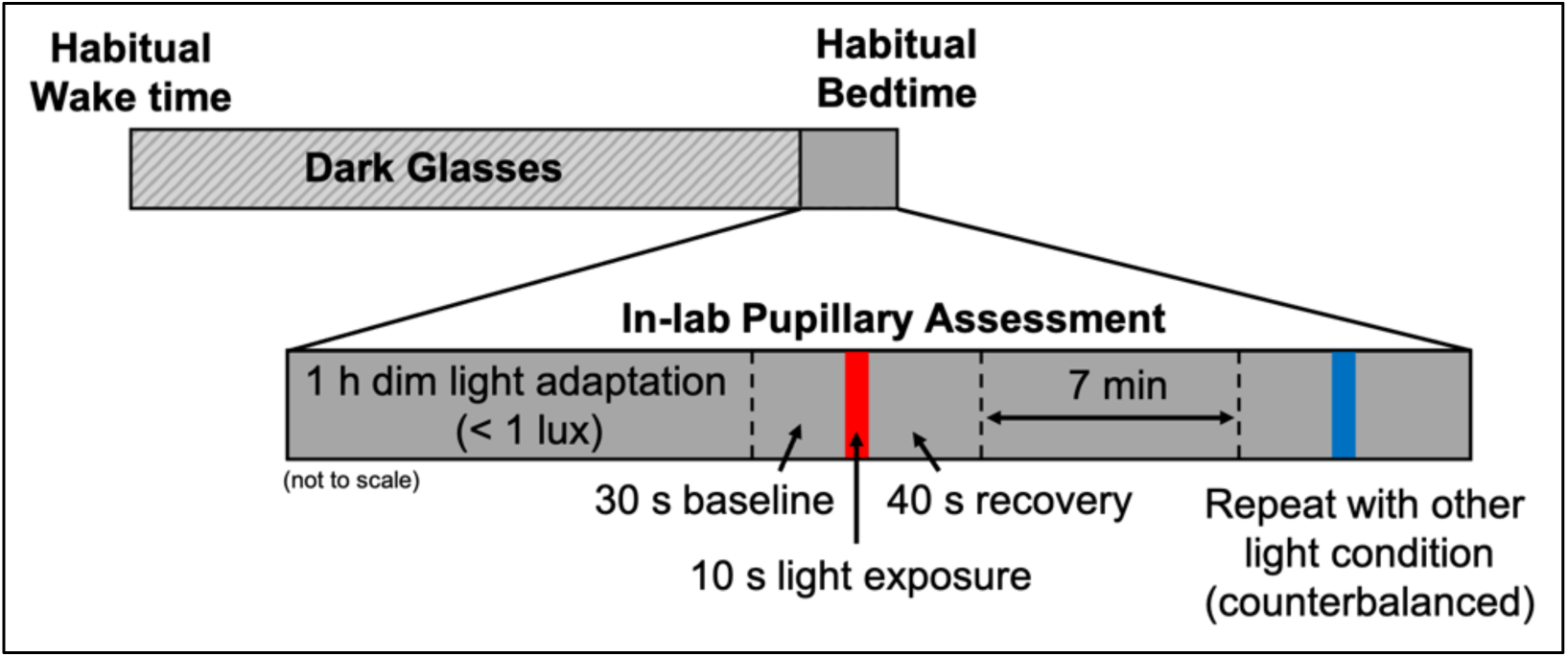
Pupillary assessment protocol. On the day of the pupillary assessment, participants remained indoors wearing a pair of dark tinted glasses from their habitual wake time until the start of the dim light adaptation at the laboratory (2 h before scheduled bedtime). Following a 1 h dim-light adaptation, pupil diameter was measured during a 30 s baseline, 10 s light exposure, and 40 s recovery. After a 7 min dim-light re-adaptation, the measurement was repeated for the remaining light condition, counterbalanced across participants.

The pupillary assessment then began 1 h before each participant’s scheduled bedtime. Researchers fitted the participant with a head-mounted eye tracking unit (ETL-100F, ISCAN, Inc., Woburn, MA, USA), which measures pupil diameter at a rate of 60 Hz. Participants were seated at a table and instructed to keep their chin on a chinrest ∼42 cm from the light source and look straight ahead during all measurements. Pupil diameter was recorded during a 30 s baseline in darkness, a 10 s light stimulus, and a 40 s recovery period in darkness to measure pupillary re-dilation. Participants were exposed to either a blue or red narrow-band LED light source matched in photon flux at 3.0×10^13^ photons/cm^2^/s (**Table 1; Fig. S2**). The light source consisted of LED strips (YUNBO, Changzhou, China) arranged on a backplane of a 61 cm × 61 cm × 12 cm deep white box with an acrylic diffusing panel at the front. A research spectrometer (MSC15, Gigahertz-Optik, Amesbury, MA, USA; calibrated 04/01/2021) was used to verify light intensity on the vertical plane from the chinrest before each participant. Following a 7 min re-adaptation to the dim-light, the measurement was repeated for the remaining lighting condition, the order of which was counterbalanced across participants. For four participants (three children, one adolescent), a visual scan of the data revealed an error in the recording. These children were re-adapted to the dim light for an additional 7 min before the measurement was repeated. Light metrics are reported in accordance with the ENLIGHT Checklist and Guidelines^31^.

**Table 1.** Spectral characteristics for each experimental light source. Measurements were obtained with a research spectrometer before each participant’s assessment. Means (standard deviations) across all participants are presented in the table below.

| **Light parameter** | **Blue** | **Red** |
| --- | --- | --- |
| **Peak wavelength (nm)** | 459.08 (0.36) | 627.12 (0.29) |
| **Full-width half maximum (nm)** | 24.56 (0.15) | 20.67 (0.13) |
| **Photopic illuminance (lux)** | 7.17 (0.24) | 22.15 (0.67) |
| **Melanopic equivalent daylight illuminance (lux)** | 70.09 (2.32) | 0.10 (0.01) |

### Analysis

Actigraphy data were scored using our standard, published methods^32^. Light exposure at the wrist was recorded by the actigraph in 1 min epochs. Light history across the first five days of the protocol was calculated as the average across each epoch between 00:00 on Day 1 and 23:59 on Day 5 of the protocol. Light history on Day 6 was calculated as the average across epochs between 00:00 and each participant’s arrival at the lab for their pupillary assessment (2.5 h before their scheduled bedtime), at which time the actigraph was removed and each participant was exposed to the same lighting environment in the lab prior to the PLR measurement.

Pupil diameter data were analyzed only from the right eye of each participant and recorded in raw pixels. All data processing was performed using Matlab R2021b (The MathWorks, Inc., Natick, MA, USA). Prior to extracting variables, the data were filtered and smoothed to remove blinks. This was achieved by first manually setting upper and lower thresholds to exclude outliers due to blinks. Then, the data were smoothed using the Matlab ‘lowess’ smooth function with a span of 0.01. The PLR variables (adapted from Najjar et al. 2021^12^) were then extracted from this smoothed data (**Table 2)**. First, a baseline pupil size, in pixels, was calculated as the median diameter across the 30 s prior to light onset. Each data set was then normalized to this baseline to translate the pupil size in pixels to percent constriction away from baseline, with 0% representing baseline pupil size.

**Table 2.** Description of each pupillary variable calculated.

| **Variable Name (units)** | **Definition** |
| --- | --- |
| Baseline Pupil Diameter (pixels) | Median pupil diameter in the 30 s preceding light onset |
| Phasic Constriction<br>(% from baseline) | Median % constriction between 300-700 ms after light onset |
| Constriction Latency (s) | Time required after light onset for the pupil to reach 10% of constriction |
| Maximum Pupil Constriction<br>(% from baseline) | $1 - \frac{\text{Min Pupil Diameter (during light on)}}{\text{Baseline Pupil Diameter}}$ |
| Sustained Slope (%/s) | Slope of best fit linear regression from 3-10 s after light onset |
| Post-Illumination Pupil Response<br>(PIPR, % from baseline) | Median percent constriction between 5 to 7 s after light offset |
| Area Under the Curve<br>(AUC, %.s) | Absolute value of a trapezoidal numerical integral of the data between light offset and 12 s after light offset. |

Phasic constriction and constriction latency could not be determined for two children due to a technical error in recording the exact moment of light onset during one of their measurements.

All statistical analyses were conducted with SPSS Statistics 28.0 (IBM Corp. Armonk, NY, USA). Due to non-normal distributions, Mann-Whitney U tests were used to examine group differences in light history. A Wilcoxon signed rank test was used to examine individual changes in light history between Days 1 through 5 compared with Day 6. An independent samples t-test was utilized to examine group differences in baseline pupil diameter. PLR data were analyzed with mixed-model ANOVAs to examine differences in response to the two light conditions (Red, Blue) and across the age groups (Children, Adolescents). Effect sizes for significant results are provided as partial-eta squared. Post-hoc comparisons were performed with repeated measures t-tests, and effect sizes were determined using Cohen’s d. An alpha-level of 0.05 was used to determine statistical significance.

## RESULTS

Two participants were myopic, but did not wear corrective lenses during the pupillary assessments. Analyses were performed both including and excluding their data. As expected from the previous literature^33^, we observed no differences in the results of the analyses after excluding these participants.

### Prior light exposure

Across the first five days of the protocol, average light exposure as measured from the wrist was significantly greater for children (M = 248.93 lux, SD = 80.88) than adolescents (M = 162.08 lux, SD = 147.94; z = -2.99, p < 0.01, r = 0.47). On the day of the pupillary assessment (Day 6), however, when participants were instructed to remain indoors throughout the day, no difference in light exposure between the two age groups was observed (children: M = 91.23 lux, SD = 70.28, adolescents: M = 64.82 lux, SD = 52.20; z = -1.26, p = 0.22). Additionally, across individual participants, average light exposure on Day 6 was significantly less than on Days 1 through 5 (z = -4.80, p < 0.001, r = 0.76).

### Baseline pupil size

Baseline pupil diameter before each participant’s first light exposure was not significantly different between children (M = 231.62 pixels, SD = 35.31) and adolescents (M = 242.88 pixels, SD = 29.23; t(38) = -1.09, p = 0.28). Neither was any group difference observed in baseline pupil diameter before each participant’s second light exposure (t(38) = -1.69, p = 0.10; M = 225.49 pixels, SD = 34.42 for children; M = 242.18 pixels, SD = 27.31 for adolescents). Descriptive statistics for each PLR variable across light color and age group are presented in **Table S1.**

### Features of pupillary light constriction

Average pupil constriction (as a % away from baseline) was plotted for each age group and light condition (Figure 2). Across both age groups, phasic constriction was marginally greater in response to blue than red light (p = 0.054, η_p_^2^ = 0.10; **Fig. 3A**). Regardless of the wavelength of the stimulation, children had both a significantly greater phasic constriction (p < 0.01, η_p_^2^ = 0.18) and smaller constriction latency (p = 0.01, η_p_^2^ = 0.17; **Fig. 3B**) than adolescents, indicating a faster and more robust initial pupillary response to light in general. Maximum pupil constriction was greater in response to blue light compared to red light for both age groups (p < 0.01, η_p_^2^ = 0.48; **Fig. 3C**). Children also exhibited a greater maximum constriction than adolescents (p < 0.01, η_p_^2^ = 0.22). Between 3 to 10 s after light onset, pupil constriction was more sustained (i.e., a smaller magnitude slope was observed) during blue compared to red light across both age groups (p < 0.01, η_p_^2^ = 0.41; **Fig. 3D**). Conversely, an independent samples t-test revealed a larger magnitude slope during blue light in children than in adolescents (t(38) = 2.32, p = 0.03, d = 0.74), indicating a less sustained response.

**Fig. 2.**
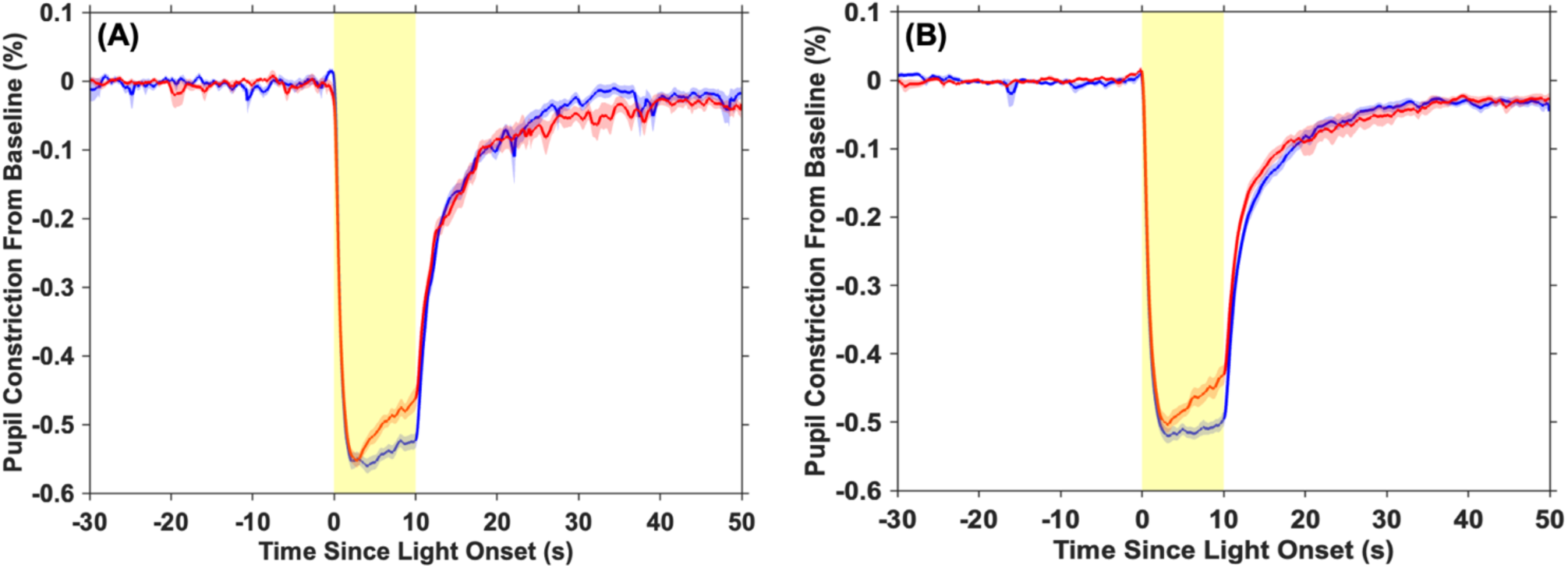
Average pupil constriction adjusted to baseline pupil diameter across the 80 s measurement. Average pupil constriction is displayed at each time point for **(A)** children and **(B)** adolescents for red and blue light separately. The shaded red and blue areas around each average line represent the standard error of the mean. The shaded yellow area denotes the timing of the 10 s light exposure.

**Fig. 3.**
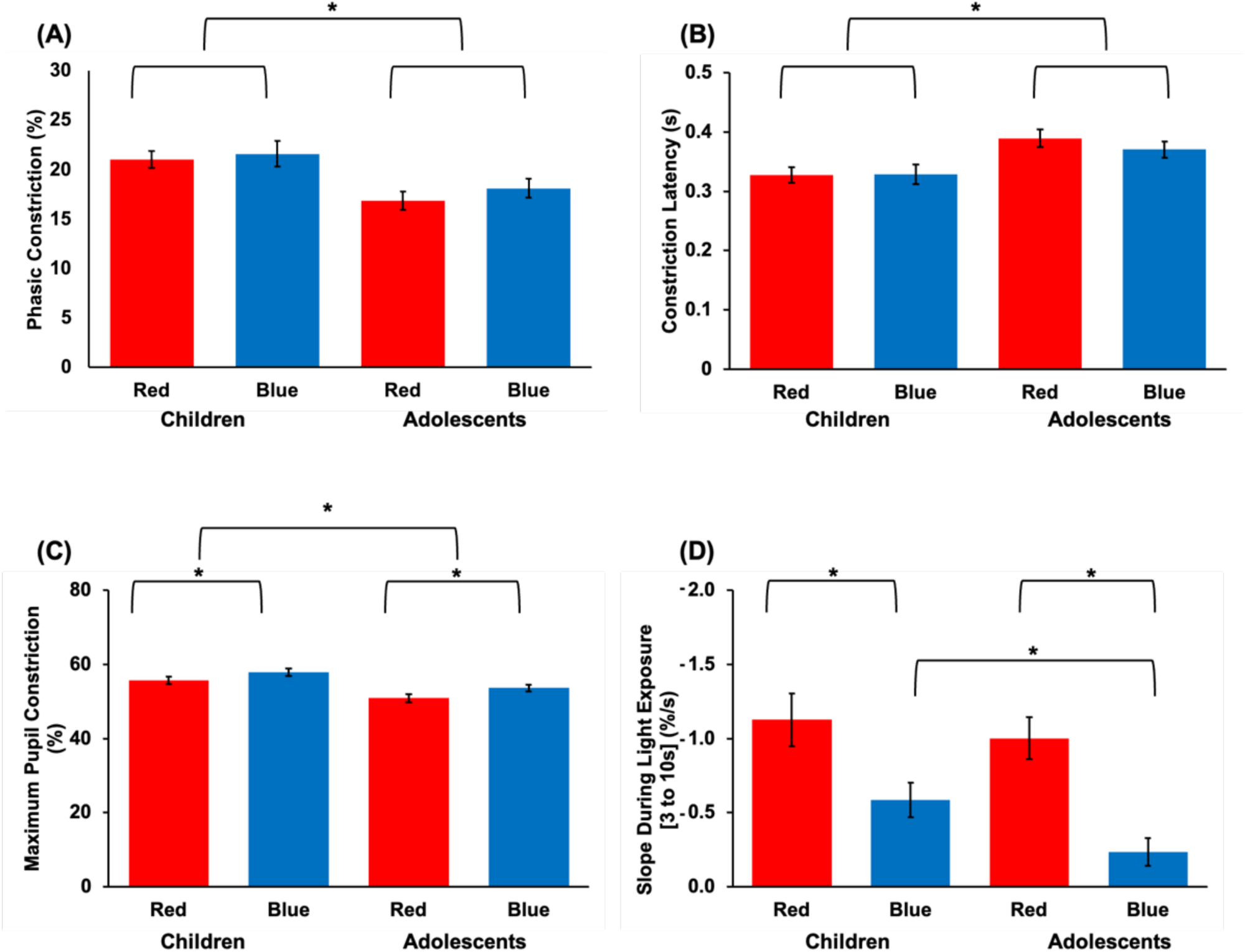
PLR metrics during light onset. Results depicting **(A)** phasic constriction; **(B)** constriction latency; **(C)** maximum pupil constriction; and **(D)** slope during light exposure broken down by age group and experimental light condition. Asterisks denote statistical significance (p < 0.05).

### Features of pupillary re-dilation

The area under the curve (AUC) 0-12 s after light offset was significantly larger for blue compared to red light (p = 0.045, η_p_^2^ = 0.10; **Fig. 4A**). However, post-hoc t-tests revealed that this was observed for adolescents (p < 0.01, d = 0.72), but not children (p = 0.66). Similarly, the PIPR was larger following blue compared to red light in adolescents (p = 0.02, d = 0.60; **Fig. 4B**), but not in children (p = 0.76).

**Fig. 4.**
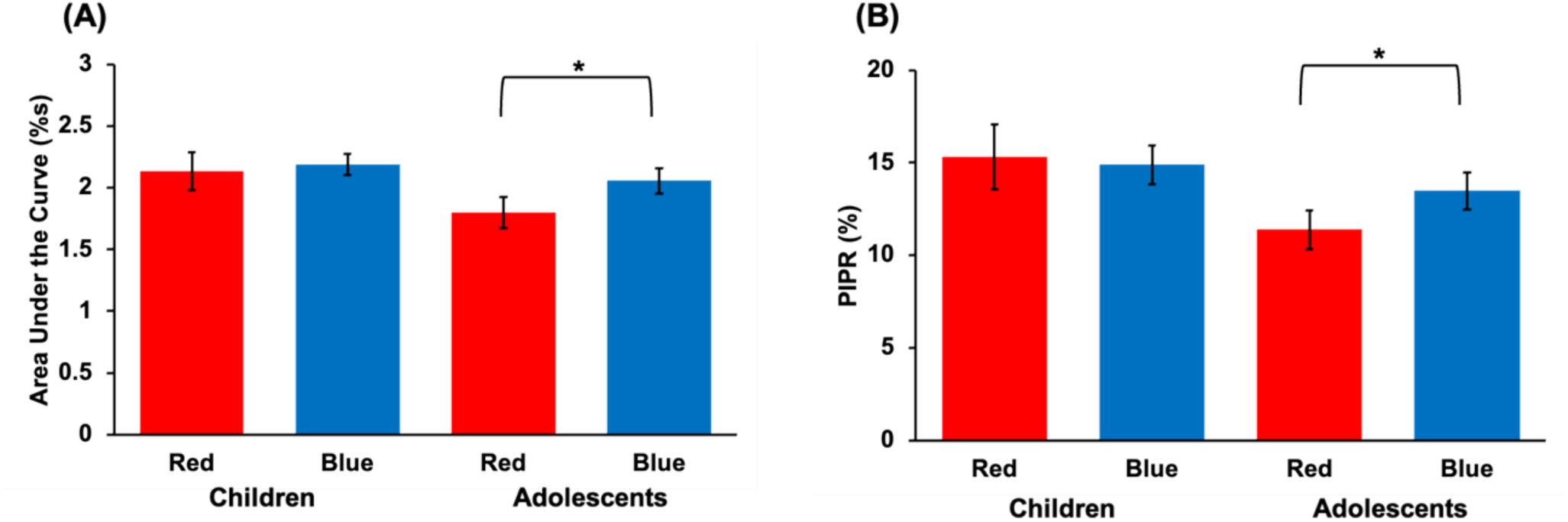
PLR metrics following light offset. Results depicting **(A)** area under the curve and **(B)** post-illumination pupil response (PIPR) broken down by age group and experimental light condition. Asterisks denote statistical significance (p < 0.05).

## DISCUSSION

In this well-controlled, within-subject study examining age-related changes in the pupillary light reflex, we observed a greater and more sustained pupillary response to blue compared to red light presented in the hour before habitual bedtime for both school-aged children and adolescents. Additionally, although children exhibited a faster and greater amplitude of transient and maximal pupillary constriction across both light colors compared to adolescents, they yielded a reduced sustained response to blue light.

Previously, Crippa and colleagues ^23^ reported that children aged 3-10 years required a brighter red light stimulus to reach 5% pupil constriction than children 10-18 years. Additionally, in children aged 6-17 years, pupillary constriction in response to a green light stimulus has been shown to increase until age 8 before plateauing ^16^. Similarly, latency was observed to decrease across ages 6-9, then stabilized. In contrast, our findings suggest that maximum constriction to both red and blue light decreases and latency increases again between school-age and adolescence. By examining more narrow age groups (8-9 and 15-16 years), our data offer a more fine-grained examination of age-related differences in the PLR than prior published findings, with particular focus around the onset of puberty where changes in the non-visual response to light have been reported ^34^.

Our findings contribute to a growing literature demonstrating maturation of the non-visual system throughout development. Previous work by Crowley, et al.^34^ demonstrated a reduced circadian response to light in later adolescence. Pre-pubertal adolescents exhibited significantly more melatonin suppression to evening light exposure than post-pubertal adolescents. Additionally, school-aged children display nearly twice as much melatonin suppression to evening bright light as adults^20^. Our data are also in agreement with experimental findings in mice showing that the pupillary light reflex continues to develop until adulthood. In that study, the PLR in response to blue and red light was compared in 1-month-old mice to 2-month-old mice and adult 4-month-old mice. The 1-month-old mice exhibited greater maximal constriction to both light stimuli than 2-month-old mice, as well as a smaller sustained constriction response to blue light ^35^, strongly mirroring our present findings in humans.

The post-illumination pupil response, especially 1.7 s after light offset, is reported to be entirely driven by melanopsin^8^. Consistent with previous findings in adults^25, 36^, post-illumination metrics from our study (i.e., AUC and PIPR) were greater in response to blue compared to red light in adolescents. This difference, however, was not observed in our sample of school-aged children, although the amplitude of the post-illumination response was at least as large as that of the adolescents. Coupled with the children’s smaller sustained response to blue light compared with adolescents, these findings could suggest continued development of the melanopsin-driven pupillary response to evening light during youth.

Our data were collected utilizing a rigorous protocol designed to control for several confounding variables that could influence the pupillary light reflex. By remaining indoors and wearing a pair of dark glasses throughout Day 6 of the protocol, participants were exposed to significantly less light on the day of the assessment, with less individual and group variability in light history.

Additionally, participants restricted light exposure between bedtime and wake time throughout the protocol to provide consistent light/dark timing. Finally, the pupillary assessment began 1 h before each participant’s habitual bedtime in order to anchor the assessment within a consistent circadian window across individuals. By focusing only on the hour before bedtime, our findings offer a more translatable examination of how the light exposure children and adolescents receive before bedtime stimulates the non-visual system, potentially leading to circadian disruption.

Our study has a few limitations. The phase angle between melatonin onset (a marker of the timing of the circadian clock) and bedtime grows wider across adolescence ^37^. Because we anchored the protocol to habitual bedtime and did not assess melatonin onset, the exact circadian timing of the pupillary assessment could have differed between the age groups. Furthermore, adherence to wearing the dark glasses on the final day of the protocol was confirmed by self- or parental-report and could not be verified objectively. Finally, participants in our sample were predominantly reported as Caucasian by their parents (87.5%). Previous studies with children and adults reported racial differences in the pupillary light reflex and circadian sensitivity to light ^38, 39^. Therefore, given the lack of diversity in our sample, our findings may not be representative of the PLR in children and adolescents across different racial and ethnic identities.

In conclusion, in this well-controlled, within subject study, we highlight differences in the pupillary response to blue and red light between children (8-9 years) and adolescents (15-16 years). While our findings demonstrate an overall heightened sensitivity to evening light in children compared to adolescents, they also suggest continued maturation of the human non-visual photoreception/system throughout development.

## ACKNOWLEDGEMENTS

We thank the participants and families who took part in this research. We also thank the students and staff of the Sleep and Development Lab at the University of Colorado Boulder who assisted in data collection.

