## Supplemental Materials for "Differences in the Pupillary Responses to Evening Light between Children and Adolescents"

**Fig. S1. Visible light transmission through dark glasses.**

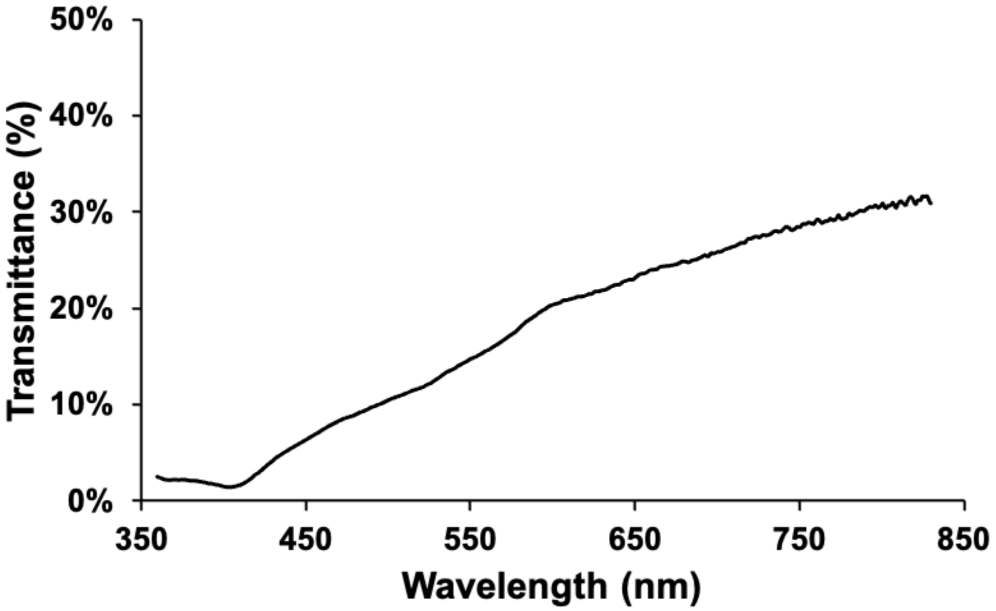

**Fig. S2. Spectra of experimental light conditions.**

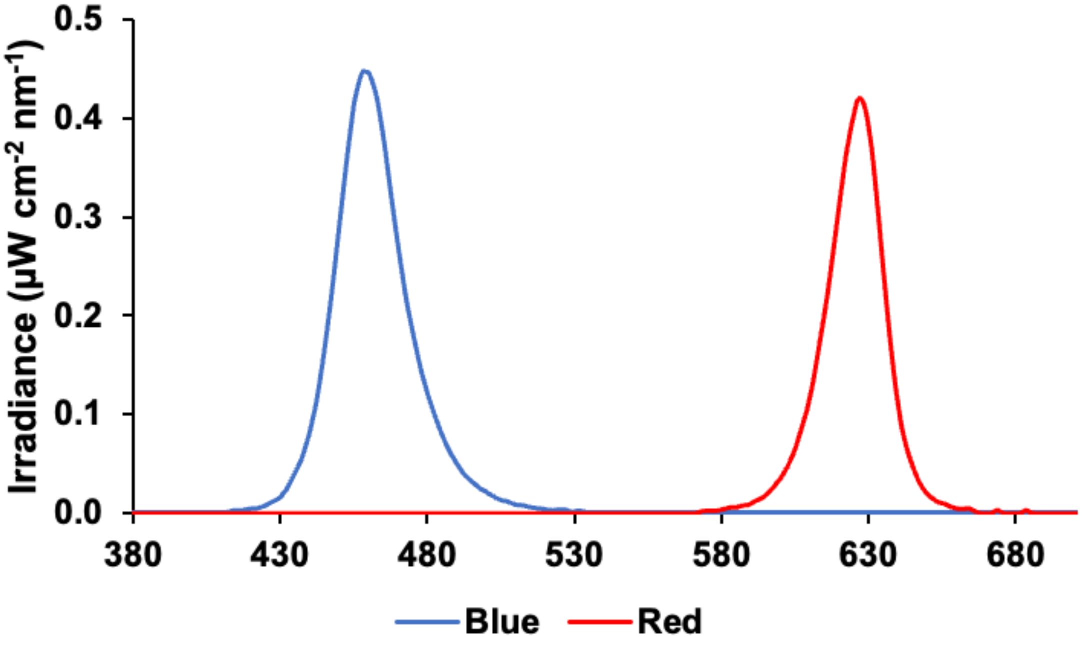

**Table S1**. Means and standard deviations of each pupillary feature broken down by age group and experimental light condition.

| Variable | Children | | Adolescents | |
|---|---|---|---|---|
| | Red | Blue | Red | Blue |
| Phasic Constriction (%) | 21.17 (0.85) | 22.51 (1.32) | 16.86 (0.94) | 18.13 (0.95) |
| Constriction Latency (s) | 0.33 (0.01) | 0.33 (0.02) | 0.39 (0.01) | 0.37 (0.01) |
| Max Constriction (%) | 55.72 (0.99) | 57.85 (1.03) | 50.87 (1.15) | 53.62 (0.93) |
| Sustained Slope (%/s) | -1.13 (0.18) | -0.59 (0.12) | -1.00 (0.14) | -0.24 (0.09) |
| PIPR (%) | 15.31 (1.74) | 14.89 (1.05) | 11.39 (1.05) | 13.48 (1.02) |
| AUC (%.s) | 2.13 (0.15) | 2.19 (0.09) | 1.80 (0.12) | 2.06 (0.11) |
